# Nanoscale architecture and coordination of actin cores within the sealing zone of human osteoclasts

**DOI:** 10.1101/2021.12.09.471901

**Authors:** Marion Portes, Thomas Mangeat, Natacha Escallier, Brigitte Raynaud-Messina, Christophe Thibault, Isabelle Maridonneau-Parini, Christel Vérollet, Renaud Poincloux

## Abstract

Osteoclasts are unique in their capacity to degrade bone tissue. To achieve this process, osteoclasts form a specific structure called the sealing zone, which creates a close contact with bone and confines the release of protons and hydrolases for bone degradation. The sealing zone is composed of actin structures called podosomes nested in a dense actin network. The organization of these actin structures inside the sealing zone at the nano scale is still unknown. Here, we combine cutting-edge microscopy methods to reveal the nanoscale architecture and dynamics of the sealing zone formed by human osteoclasts on bone surface. Random illumination microscopy allowed the identification and live imaging of densely packed actin cores within the sealing zone. A cross-correlation analysis of the fluctuations of actin content at these cores indicates that they are locally synchronized. Further examination shows that the sealing zone is composed of groups of synchronized cores linked by α-actinin1 positive filaments, and encircled by adhesion complexes. Thus, we propose that the confinement of bone degradation mediators is achieved through the coordination of islets of actin cores and not by the global coordination of all podosomal subunits forming the sealing zone.

## Introduction

Osteoclasts are giant multinucleated cells of the hematopoietic lineage, specialized in the degradation of bone matrix. To do so, protons accumulate into the resorption lacuna thanks to the vacuolar H^+^-ATPase (v-ATPase), thus lowering the local pH and facilitating the solubilization of apatite, the main mineral component of bone. This acidic environment is also prone to enhance the digestion of bone organic matrix by proteases secreted by osteoclasts such as cathepsin K ^1, 2^. The efficiency of this process relies on the ability of the cell to create an enclosed resorption compartments, *via* the formation of a unique cytoskeletal structure, the sealing zone ^3, 4^.

First described as a subcellular entity made of electron dense material and apparently deprived of any organelle, thus resulting in first denomination as the “clear zone”, the sealing zone was then revealed to consist in a dense accumulation of actin filaments forming a circular shape surrounding the resorption lacuna ^5–7^. Examination with scanning electron microscopy (SEM) of cells removed of their basal membrane brought to light the peculiar arrangement of actin filaments within this structure, and particularly unveiled the existence of a dense network of podosomes composing this structure ^8–10^. Podosomes are adhesion structures that are generally scattered in macrophages and dendritic cells. Podosome typical components, such as vinculin, paxillin, talin or cortactin, are also localized in the sealing zone ^11–19^. In a distinct way compared to single podosomes, vinculin, paxillin and talin were reported to form a “double circle” flanking the sealing zone on each side, while cortactin mainly colocalized with actin in between ^3, 13, 15, 20–22^. Noteworthy, most of the studies on the bone degradation machinery were carried out based on observations of osteoclasts on glass substrates instead of bone, due to the lack of optical transparency and high auto-fluorescence of the mineralized matrix. As a result, only poor knowledge has been collected about the architecture and dynamics of the functional sealing zone. Indeed, it has been suggested that actin structures on bone differ from the ones formed on glass, mainly in their total width, and the interconnectivity and density of actin cores within ^8^. Therefore, only structures formed on bone or bone-mimicking materials are called sealing zone, structures on glass being denominated as “sealing zone like” or podosome belts ^3, 23^.

How podosomes are organized and coordinated to allow efficient sealing and bone degradation is therefore still unknown. Hence, it appears paramount to develop higher-resolution microscopy techniques compatible with observation on bone substrates. This could yield valuable information concerning the spatial distribution of major actin-binding proteins within the sealing zone, otherwise only arduously accessible *via* electron microscopy and correlative microscopy ^8–10, 24, 25^. Additionally, observation of the sealing zone internal dynamics would provide substantial hints to understand the sealing ability of such a structure. This exploration would require both a spatial and temporal high-resolution microscopy technique.

In this work, we use cutting-edge super-resolution microscopy methods to reveal the architecture and dynamics of the bone degradation machinery formed by human osteoclasts. First, we examined the three-dimensional nanoscale organization within the podosome belt thanks to a single molecule localization method. Then, random illumination microscopy (RIM) acquisitions of human osteoclasts plated on bone allowed to resolve single actin cores composing the sealing zone. RIM technique also proved to be efficient in deciphering this nanoscale organization in living samples. Hence, cross-correlation analysis of the fluctuation of the sealing zone actin content could show that cores are locally synchronized. Further analysis of the organization of adhesion components and actin crosslinkers revealed that the sealing zone is composed of coordinated groups formed by α-actinin1-linked podosomal cores and encircled by adhesion complexes. Therefore, confinement of bone degradation enzymes may be achieved through the alternating contact of functional islets of actin cores.

## Results

### Three-dimensional nanoscale organization of the podosome belt in human osteoclasts

The sealing zone formed by bone-degrading osteoclasts is described as a dense network of actin cores that appear fused under conventional microscopy, surrounded by two lines of adhesion sites ^3, 13, 15, 20–22^. The precise architectural and dynamics of this structure are almost uncharacterized. To address the organization of osteoclast podosomes, a 3D super-resolution method called DONALD was used. DONALD is a single molecule localization method combining direct stochastic optical reconstruction microscopy for the in-plane detection of proteins, and SAF analysis to gain access to the absolute axial position of fluorophores relative to the glass coverslip. This nanoscopy technique thus benefits from approximately 15 nm localization precision in the three dimensions (3D) ^26^. Human osteoclasts derived from blood monocytes were differentiated for 10 days, plated on glass and observed with this 3D super-resolution technique. On glass, osteoclasts form structures called podosome belts, sharing certain characteristics with sealing zones. In particular, podosome belts also exhibit areas where podosome cores appear fused, *i.e*. when sets of several cores are brought together inside a single and large ring of adhesion sites, similar to the described structure of the sealing zone. Indeed, this fusion of podosome cores was observed in 9.4+/-3.5% of multinucleated human osteoclasts. We therefore focused on these fusion zones.

Spatial distributions of key structural components of podosomes, namely cortactin, α-actinin1, filamin A, vinculin, paxillin and the C-terminal extremity of talin (talin-C), whose height was previously correlated with podosome protrusion forces ^27^, were thus explored using DONALD. Image analysis consisted in localizing the various target proteins with respect to the actin cores composing the podosome belts (Figure S1). Cortactin in-plane distribution was characterized by an accumulation in the first 250 nm surrounding the actin cores, and kept at a rather constant height of approximately 164 nm (Figure S2A-A’’’). α-actinin1 was preferentially localized in the vicinity of the actin cores, up to 500 nm in distance. Similarly to cortactin, its height stayed approximately constant throughout the distance profile, at approximately 126 nm (Figure S2B-B’’’).

Vinculin, paxillin, talin and filamin A were mostly absent from the regions of dense actin staining, and encircled multiple actin cores (Figure 1A-D’, Figure S2C-C’’ and D-D’’). Vinculin and paxillin displayed rather similar in-plane distributions, they are 350 nm and 510 nm from actin cores on the external side of the podosome belt, and 570 nm and 510 nm from the core on the internal side, respectively (measurements not shown). In contrast, talin and filamin A displayed large ranges of preferential localization, they are situated at 630 nm for both proteins on the external side, and at 610 nm and 710 nm towards the inner part of the belt, respectively. Talin-C and vinculin heights declined by nearly 20 nm towards the inner part of the cell (Figure 1E-I). In contrast, paxillin and filamin A, which average heights were 45 nm and 139 nm, respectively, exhibited almost no change in height when away from the belt center (Figure S2C-D’’’).

**Figure 1.**
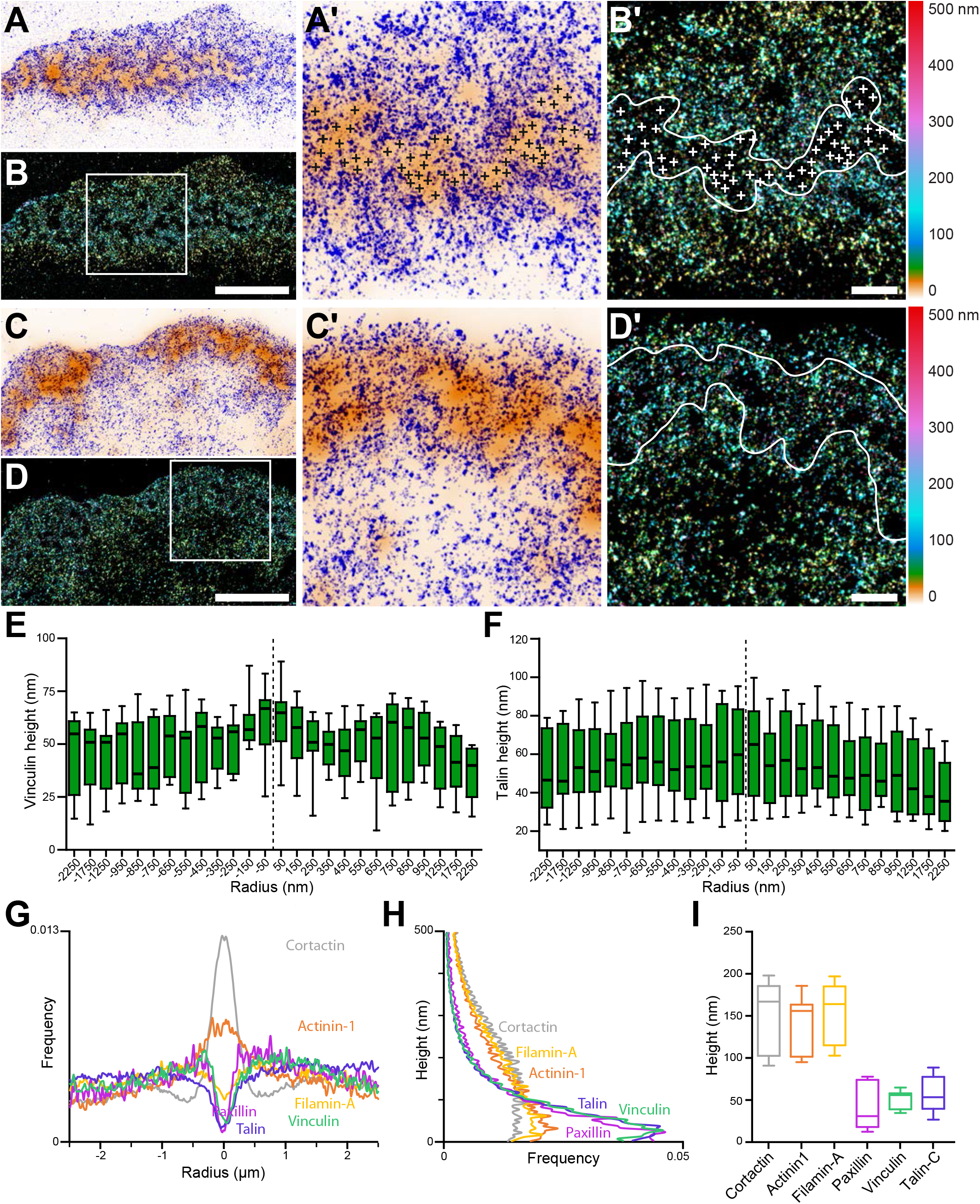
3D nanoscopy of vinculin and talin-C in the osteoclast podosome belt. (A) Representative dSTORM images of vinculin (blue) merged with the corresponding epifluorescence images of the F-actin cores (ochre). (A’) Enlarged view of (A). The black crosses indicate the localization of actin cores. (B) DONALD images corresponding to (A) where the height is represented in false color (scale shown in (B’). (B’) Enlarged view of (B). The white crosses indicate the localization of actin cores. (C) Representative dSTORM images of talin-C (blue) merged with the corresponding epifluorescence images of the F-actin cores (ochre). (C’) Enlarged view of (C). (D) DONALD images corresponding to (A) where the height is represented in false color (scale shown in (D’). (D’) Enlarged view of (D). (E-F) Height profiles for vinculin (E) and talin-C (F) with respect to the distance to the center of the podosome belt. (G-H) Radial (G) and vertical (H) distributions of cortactin, α-actinin1, filamin A, paxillin, vinculin and talin-C in podosome belts. (I) Median axial positions of F-actin, cortactin, α-actinin1, filamin A, paxillin, vinculin and talin-C in podosome belts. Scale bars: 5 μm (B, D), 1 μm (B’, D’).

### Nanoscale organization of actin cores in the sealing zone

To further explore the architecture of the sealing zone, we then investigated the organization of podosomes in mature human osteoclasts plated on bovine bone slices. After 3 days, osteoclasts efficiently degraded bone, as shown by scanning electron microscopy (SEM) observations (Figure 2A). SEM acquisitions of unroofed cells confirmed that the sealing zones formed by human osteoclasts are composed of individual F-actin cores (Figure 2B-B’), similarly to what was shown in osteoclasts differentiated from the mouse cell line RAW 264.7 or harvested from rabbit long bones ^8, 10^. These cores were nested in a dense network of actin filaments, and appeared connected to their neighbors by filaments running parallel to the substrate (Figure 2B’’, arrowheads). The morphometric characteristics of the network were assessed by manually encircling each core and estimating their radius from the selected area. Analysis of 457 cores in 9 different cells yielded a median radius of 114 nm (Figure 2D). Furthermore, applying Delaunay’s tessellation to each dataset allowed for the characterization of the average inter-core distances within the same cell. Direct neighbor pairs were 705 nm apart (Figure 2E), and first neighbors were 443 nm apart (Figure 2F), similarly to what was described previously ^28^.

**Figure 2.**
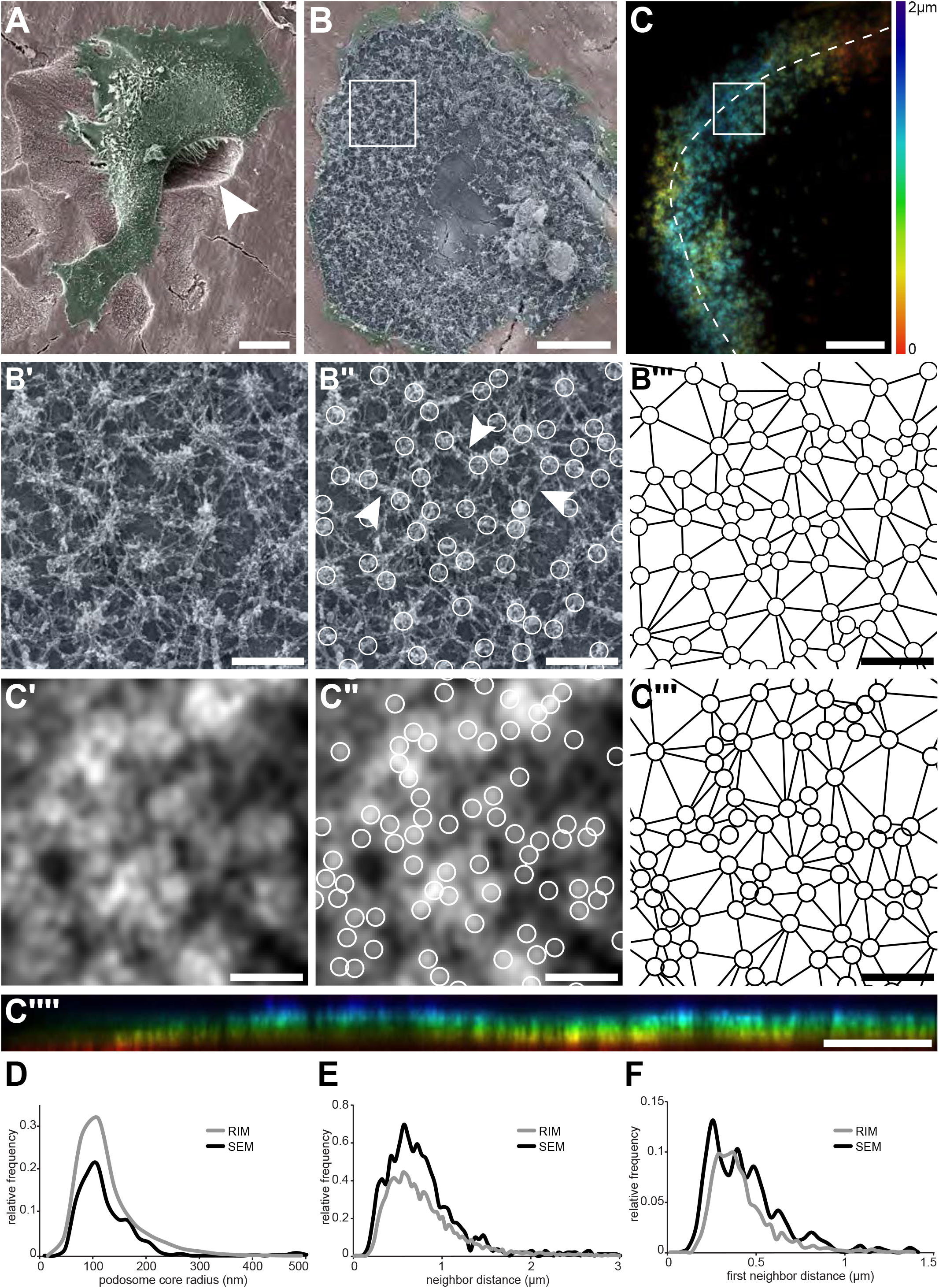
Nanoscale organization of actin cores in the sealing zones formed by human osteoclasts. (A) Pseudo-colored scanning electron micrograph of a human monocyte-derived osteoclast degrading bone. (B) Scanning electron microscopy image of an unroofed osteoclast. (B’) Enlarged view of (B). (B’’) Localization of actin cores (circles). Arrowheads point to lateral actin filaments linking actin cores together. (B’’’) Delaunay triangulation from the cores in (B’’). (C) RIM image of a sealing zone stained for F-actin. Color-coded for height using a rainbow scale. (C’) Enlarged view of (C). (C’’) Localization of actin cores (circles). (C’’’) Delaunay triangulation from the cores in (C’’). (C’’’’) Orthogonal projection along the line marked in (C) The same color code for height was used. (D) Histogram of the core radii, as measured by SEM (black) and RIM (grey). (E) Histogram of the average distances to direct neighbors measured as Delaunay edges, as measured by SEM (black) and RIM (grey). (F) Histogram of the average distances to first neighbors, as measured by SEM (black) and RIM (grey).In (D-F), 457 and 2781 cores were quantified for SEM and RIM, respectively. Scale bars: 20 μm (A), 5 μm (B, C, C’’’’), 1 μm (B’, B’’, B’’’, C’, C’’).

F-actin distribution in human osteoclasts unroofed on bone was then observed by a new super-resolution method called random illumination microscopy (RIM). RIM consists in illuminating a sample with a series of random speckles and processing the stack of images using signal processing and statistical tools, to gather a lateral and axial resolution of 100 and 300 nm, respectively. This method benefits from similar resolution as with traditional structured illumination microscopy (SIM), while not requiring any initial calibration step. In addition, its super-resolution capability allows for characterization of events within thick samples, until 30 μm in unknown optical medium ^29^. RIM thus enabled super-resolution imaging of the functional sealing zone. Actin staining showed a dense though discontinuous pattern within the sealing zone (Figure 2C,C’’’’, Figure S3, Movie1). Signal analysis localized local intensity maxima, the geometric features of which were assessed by extracting both the coordinates of the local maxima, and the signal intensity values along 8 directions, evenly distributed from the center. Signal variations were quantified along these 1 μm-long segments, the half-width of the peak in each direction was computed after spatial derivation, and the peak radii were computed by averaging the 8 values (Figure S4). The radius distribution yielded an average value of 97 nm (Figure 2D). Delaunay’s tessellation was applied to the peak coordinates to characterize their spatial arrangement. Direct neighbors were in a 694 nm distance range (Figure 2E), with first neighbors being 399 nm apart (Figure 2F).

As the distribution of radii and neighbor distances were similar with SEM and RIM, we concluded that RIM efficiently allows for the observation of single actin cores within the sealing zones of human osteoclasts.

### Local synchrony of sealing zone actin cores

We then used RIM to image living human osteoclasts adhering on bone to address the dynamics of actin cores within the sealing zone. Human osteoclasts transduced with GFP-tagged LifeAct lentiviruses and actively degrading bone slices were first observed by wide field fluorescence microscopy over 30 min. Subsequent deconvolution of the images and color-coding for time using a rainbow scale revealed that cores appeared stable over at least 30 min (Figure 3A-A’’, Movie2). Furthermore, analysis of kymographs along the sealing zone revealed actin intensity oscillations during the entire duration of the acquisitions. To further characterize this dynamic process at a higher spatial and temporal resolution, small regions of the sealing zone were observed with RIM. Variations of the actin content in podosome cores were measured and appeared to fluctuate synchronously between neighbors (Figure 3B-B’, Movie3, left).

**Figure 3.**
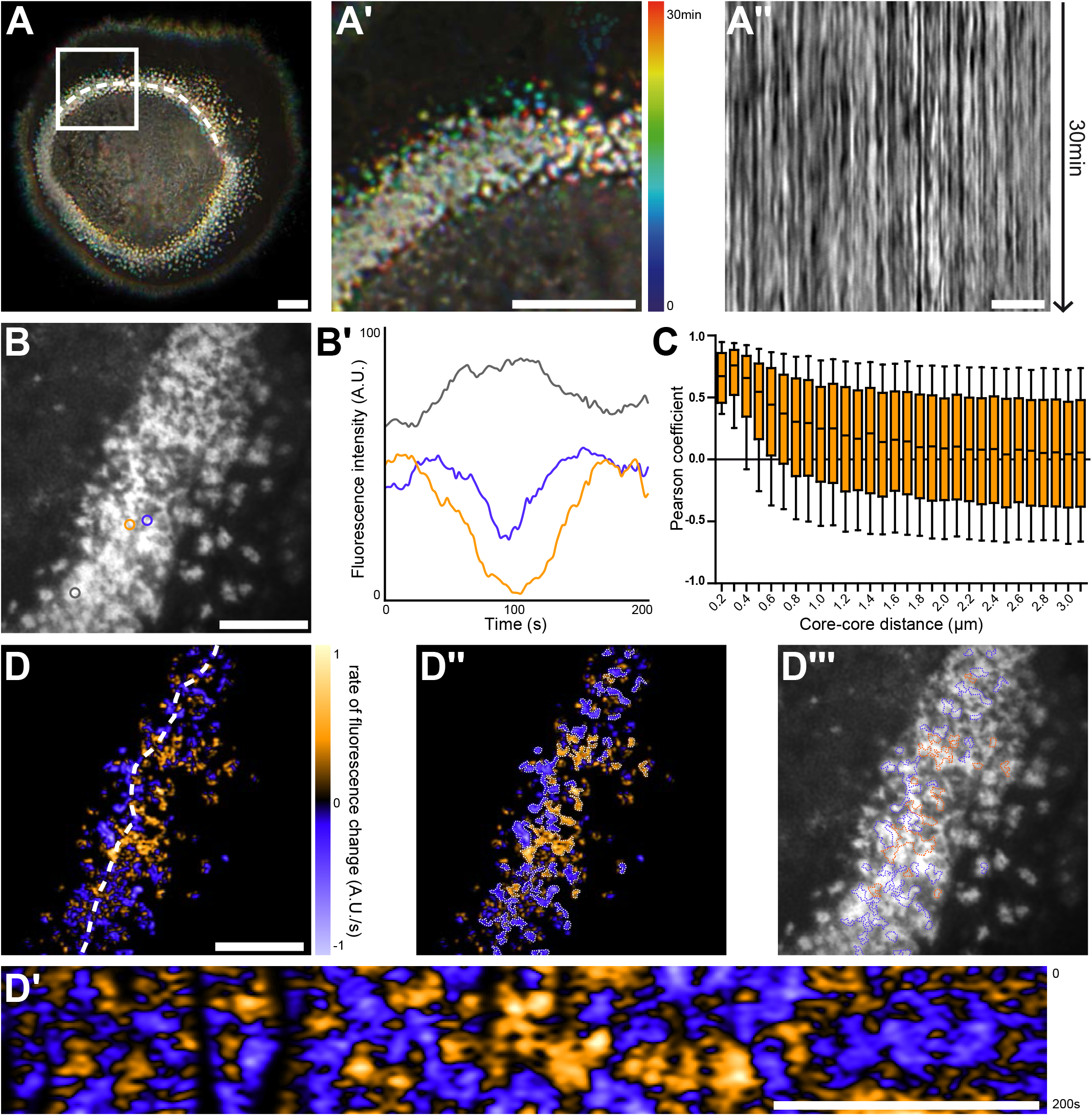
Nanoscale analysis of the dynamics of the sealing zone. (A) Temporal projection of deconvolution images of a sealing zone acquired over 30 min, color-coded for time using a rainbow scale. Thus, the structures that remain at the same spot tend to appear whiter, whereas short-lived or mobile podosomes remain colored. (A’) Enlarged view of (A). (A’’) Kymograph along the line marked in (A). (B) RIM image of a sealing zone stained for F-actin with Lifeact-GFP. (B’) Measurements of Lifeact-GFP intensity variations of actin cores marked in (B). (C) Pearson coefficients of actin intensity fluctuations of podosome pairs as a function of distance between pairs (2839 cores). (D) Image of the rate of fluorescence change corresponding to the cell shown in (B). (D’) Kymograph along the line marked in (D). (D’’) Segmentation of the growing and decreasing clusters of actin cores. (D’’’) Superimposition of the RIM image with the segmented regions of coordinated actin clusters shown in (D’’). Scale bars: 10 μm (A), 5 μm (A’-D’).

To quantify to what extent podosome actin content varied concomitantly between neighbors, 2840 cores distributed in 10 different sealing zones were localized, and their associated actin intensity signal was extracted for further analysis. The distance between every possible core pair was estimated and their respective signals were compared using Pearson cross-correlation analysis. A high positive Pearson coefficient corresponded to acute temporal synchrony for the analyzed pair of actin signals. Highest Pearson coefficient values (over 0.37) were obtained for cores within a 700 nm radius distance (corresponding to half of the maximum value). They slowly decreased to approximately 0.10 at longer distances (Figure 3C). This result hinted at the existence of a spatial synchrony between neighbors within the sealing zone, comparable to observations for podosomes in human macrophages ^30^. Moreover, analysis of the Fourier spectra corresponding to live acquisitions yielded three specific frequencies that seem to prevail: 0.01 Hz, 0.04 Hz and 0.15 Hz, corresponding to actin content fluctuation periods of approximately 100 s, 25 s and 7 s, respectively. In macrophages, similar oscillation periods had previously been identified by examining the stiffness variations of single podosomes ^31^. In order to obtain a graphical representation of podosome synchrony, differential films were assembled by subtracting 2 sequential time points, therefore representing local actin intensity gradients. Orange stands for a positive gradient, *i.e*. local polymerization, and blue represents a negative gradient, *i.e*. local depolymerization. Strikingly, polymerization and depolymerization regions appeared as clusters with slowly varying areas, containing a few actin cores (Figure 3D-D’’’’, Movie3). Actin polymerization and depolymerization processes thus appeared to be synchronous within actin core clusters.

In conclusion, we show a local synchrony of F-actin oscillations between cores of the same superstructure. This suggests the organization of actin cores into functional clusters within the sealing zone.

### Sealing zone actin cores are organized into islets surrounded by adhesion complexes

Although no partitioning of podosome cores was evident solely on the basis of the actin staining, we then reasoned that, since groups of actin cores display local synchrony, a specific organization of these cores could be revealed by localizing adhesion components of the sealing zone. Spatial distributions of the same components of the sealing zone that were imaged with the DONALD technique (Figure 1) were thus explored by RIM, and we developed a quantitative image workflow to analyze the localization of these proteins with respect to the actin cores in human osteoclasts adhering on bone. Protein localizations were evaluated along 1.5 μm long and 100 nm wide lines, in longitudinal and transverse directions relative to the local sealing zone orientation. Then, normalizing the actin core width allowed for the localization of the target proteins with respect to the core domain (see vinculin, Figure S5). Cortactin was localized within the core domain, as had been previously reported, and displayed a wider distribution compared to the average core diameter (Figure 4A-A’’,E). α-actinin1 mostly colocalized with actin at the close periphery of the core, and appeared less present in the most central part of the core (Figure 4B-B’’,E). Filamin A, vinculin, paxillin and talin were preferentially encircling multiple actin cores, with little staining in between cores in the inner part of the sealing zone (Figure 4C-C’’,E, Figure S6 and Figure S7). These observations show a strong divergence with the “double circle” distribution described before ^13, 21^, and the sealing zone appears to be composed of islets of actin cores that are bordered by a network of adhesion complexes.

**Figure 4.**
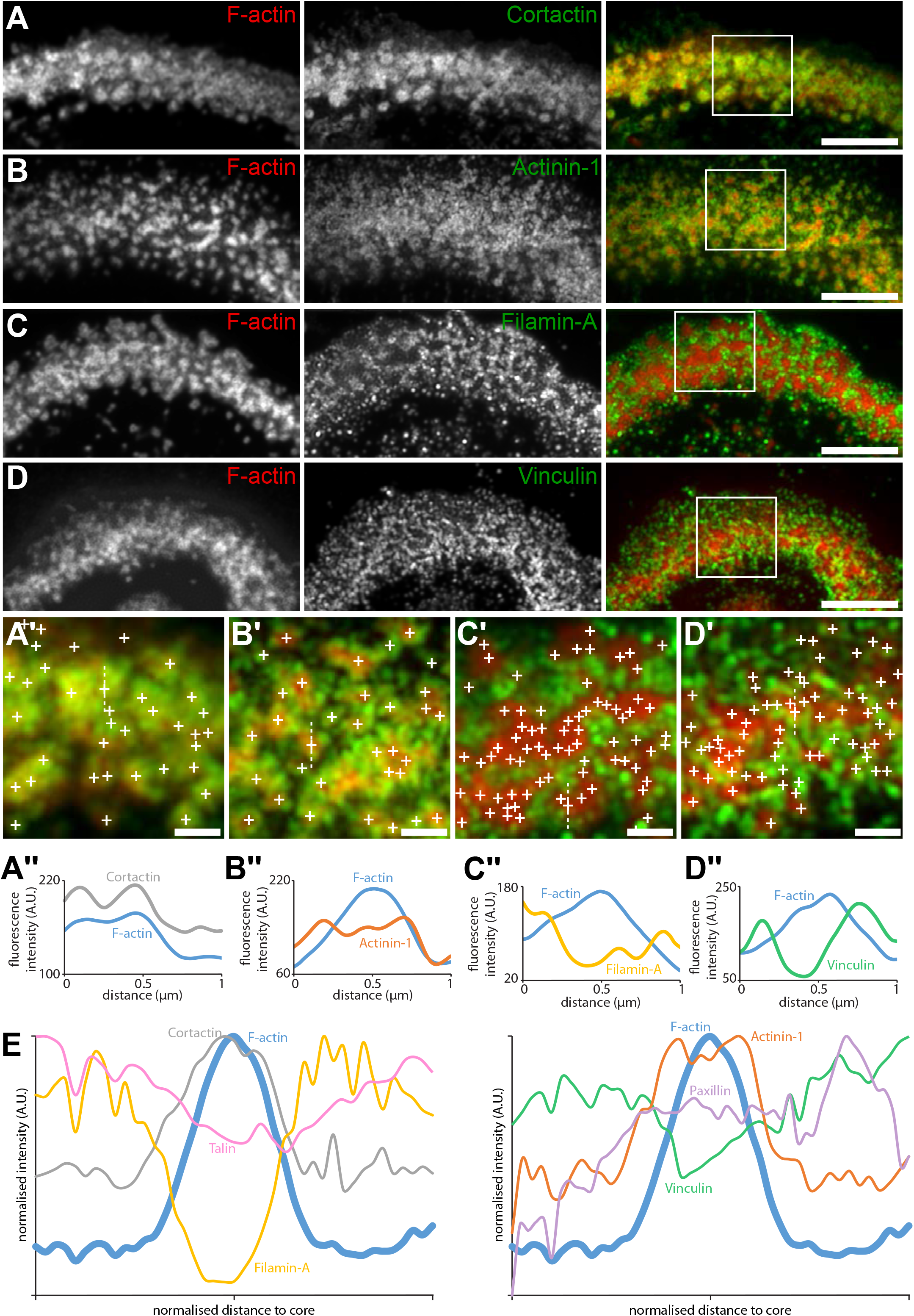
Localization in the sealing zone of actin core and ring proteins. (A) Representative immunofluorescence images of sealing zones co-stained for F-actin (red) and cortactin (green). (A’) Enlarged view of (A) where white crosses indicate the localization of actin cores. (A’’) Intensity profiles along the dotted lines marked in (A’). (B) Representative immunofluorescence images of sealing zones co-stained for F-actin (red) and α-actinin1 (green). (B’) Enlarged view of (B) where white crosses indicate the localization of actin cores. (B’’) Intensity profiles along the dotted lines marked in (B’). (C) Representative immunofluorescence images of sealing zones co-stained for F-actin (red) and filamin A (green). (C’) Enlarged view of (C) where white crosses indicate the localization of actin cores. (C’’) Intensity profiles along the dotted lines marked in (C’). (D) Representative immunofluorescence images of sealing zones co-stained for F-actin (red) and vinculin (green). (D’) Enlarged view of (D) where white crosses indicate the localization of actin cores. (D’’) Intensity profiles along the dotted lines marked in (D’). (E) Normalized intensity profiles of F-actin, cortactin, α-actinin1, filamin A, vinculin, paxillin and talin (medians of at least 200 cores for each staining). Scale bars: 5 μm (A, B, C, D), 1 μm (A’, B’, C’, D’).

In scattered podosomes formed by macrophages or dendritic cells, there is a local co-regulation between polymerization of actin in the core and formation of surrounding adhesion sites ^27, 32^. To determine whether such a local co-regulation takes place inside the islets of podosomes in the sealing zone, we first measured local intensities of cortactin, α-actinin1, filamin A, vinculin, paxillin and talin, relatively to the intensity of F-actin staining of each podosome core. These analyses revealed a positive correlation between the actin content and each of the podosome components (Figure 5A). Time-lapse RIM imaging of both F-actin and paxillin further confirmed that paxillin intensity fluctuations are locally correlated with F-actin intensity (Figure 5B-C, Movie4). These results suggest that, in addition to the local core coordination, there is also a local regulation between polymerization of actin in the core and accumulation of adhesion complex proteins.

**Figure 5.**
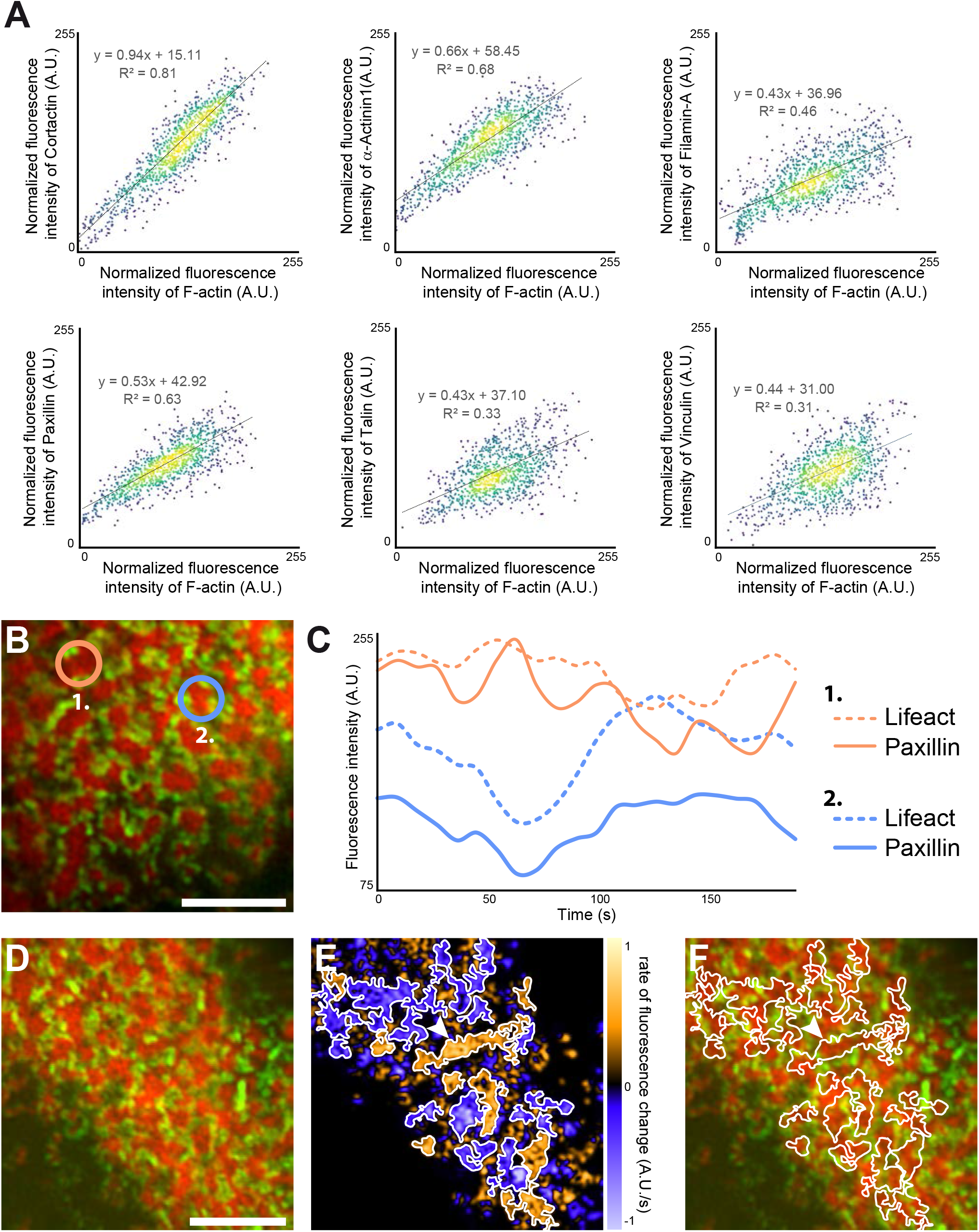
Quantification of the dynamics of the sealing zone. (A) Osteoclasts adhering to bone were stained for both F-actin and cortactin, α-actinin1, filamin A, vinculin, talin or vinculin, respectively. The intensity of each fluorescent marker in 1 μm radius circles around F-actin cores were quantified for at least 1000 cores (in five cells from different donors), and correlated to the fluorescence intensity of F-actin. Data were normalized with respect to the maximum intensity. (B-C) Time-lapse RIM imaging of F-actin and paxillin in a living osteoclast expressing Lifeact-mCh and paxillin-GFP and adhering to bone. The intensity variations of Lifeact-mCh and paxillin-GFP from two cores are shown. Note that the variations of the two podosome markers are correlated locally, but that the two cores, which are 5 μm apart, are not synchronized (C). (D) Time-lapse RIM imaging of F-actin and paxillin in a living osteoclast expressing Lifeact-mCh and paxillin-GFP and adhering to bone. A single RIM image of paxillin was acquired, followed by a stream acquisition of Lifeact-GFP, for a higher temporal resolution. (E) Image of the rate of fluorescence change corresponding to the cell shown in (D). (F) The superposition of the fluorescence image and the segmentation of clusters of synchronized areas in the sealing zone shows that there is a synchrony within zones corresponding to multiple cores encircled by paxillin. The arrowheads in (E-F) indicate a large cluster of actin cores. Note that all the different cores in this group are synchronized (E). Scale bars: 5 μm.

Finally, we explored whether actin cores observed in the same cluster surrounded by adhesion complexes were synchronous. For this purpose, we analyzed how F-actin content within cores fluctuates relatively to each other inside the same clusters. The superposition of the fluorescence image and the segmentation of clusters of synchronized areas revealed that there is a synchrony within zones corresponding to multiple cores (Figure 5D-F, Movie5).

Altogether, these data demonstrate that the actin cores which are localized within the same islets are synchronized with each other and with the surrounding adhesion complexes.

## Discussion

This study provides a nanoscale picture of the architecture, spatial organization and dynamics of podosomes in the sealing zone of human osteoclasts adhering on bone. First, we give a detailed insight into the inner organization of the sealing zone with the 3D localization of major actin crosslinkers inside the podosome belt, and examination of their axial localization hinted at the possible existence of tension within the integrin adhesion sites. Second, using RIM, we show that podosome cores were within the submicron range from their direct neighbors. Third, by monitoring podosomes in the sealing zone by time-lapse super-resolution microscopy, we characterized the long-term spatial stability of actin cores during bone resorption and we revealed a synchronous behavior for actin cores within a distance of 1 μm. Finally, we find that synchronous neighbors were grouped within clusters, which corresponded to islets surrounded by adhesion complexes composed of vinculin, talin, paxillin and filamin A. Overall these results allow for a new model of the internal architecture, dynamics and functioning of the sealing zone (Figure 6).

**Figure 6.**
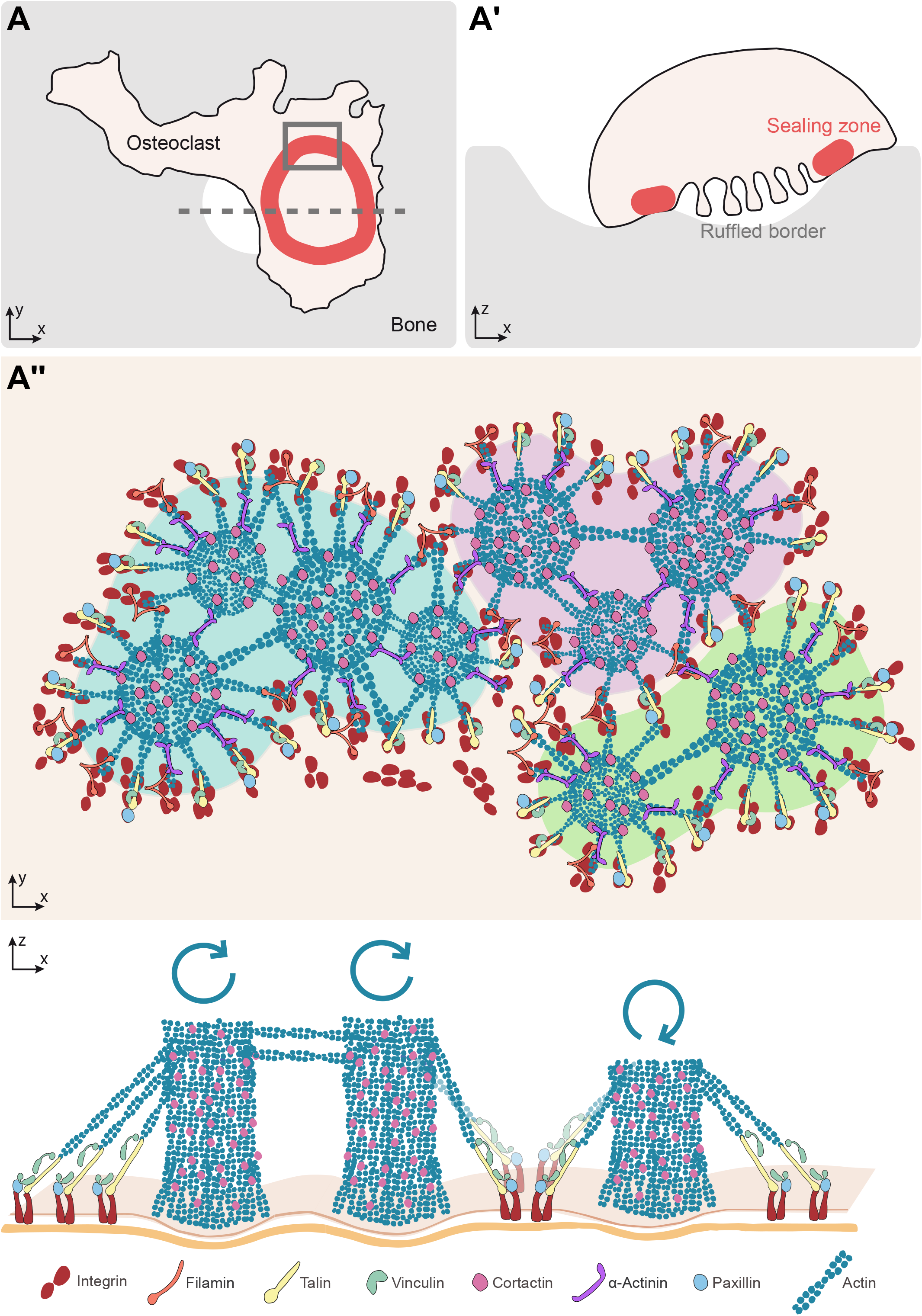
Model of the organization of the sealing zone into islets. (A-A’) Osteoclast form an actin rich superstructure called the sealing zone, in order to confine bone degradation. (A’’) Zooming inside the sealing zone reveals the 3D organization into islets of coordinated actin cores. Moreover, actin cores that are localized within the same cluster tend to display synchronized actin fluctuations, whereas there does not seem to be much correlation between clusters.

The evaluation of the 3D distributions of vinculin, paxillin, talin, filamin A, cortactin and α-actinin1 was carried out in the podosome belts of human osteoclasts on glass. Acquisitions were performed with the DONALD imaging technique, which combines dSTORM and SAF analysis for the efficient 3D detection of single fluorophores with a precision of 15 nm ^26^. This 3D nanoscopy technique has recently been applied in the context of human macrophage podosomes. It allowed for the identification of a close relationship between paxillin, vinculin and talin, and its requirement for efficient protrusion force generation ^27^. In osteoclast podosome belts, vinculin, paxillin and talin-C were localized in the close vicinity of the ventral membrane. Moreover, vinculin and talin-C appeared to rise when situated closer to the actin cores. This interesting finding could be the first insight into the identification of possible tension within the podosome belt. In fact, in podosome rings of macrophages, it was observed that when talin is stretched, vinculin height is higher, probably because vinculin binding sites are appearing when talin acquires an extended conformation ^27^. Our results with this 3D super-resolution technique were obtained for podosomes belts of osteoclasts plated on glass. While only structures formed on bone or bone-mimicking substrate are called sealing zone ^3, 23^, podosome belts share some characteristics with sealing zones such as the local fusion of podosome cores. The use of RIM allows for the identification of single actin cores within the sealing zones of human osteoclasts adhering on bone and resorbing. Podosome cores were within the submicron range from their direct neighbors, as already assessed ^28^. Additionally, we show that the sealing zone are composed of islets composed of grouped podosomes within clusters and bordered by a network of adhesion sites. These observations greatly contrasted with the “double circle” distribution (*i.e*. a dense network of actin cores that appear fused surrounded by two lines of adhesion sites) described in earlier works using conventional or confocal microscopy ^13, 21^.

Importantly, benefitting from RIM low toxicity and high temporal resolution, we could picture the sealing zone using time-lapse super-resolution microscopy. The actin dynamics was monitored within the sealing zone of osteoclasts plated on bone slices. Preliminary deconvoluted movies had revealed the apparent spatial stability of actin cores for long time periods. Actin signal processing yielded a time periodicity for these oscillations of approximately 100 seconds, which is much slower than the actin oscillations observed in scattered macrophage podosomes ^27^. Here, local oscillations associated with actin remodeling within the cores were revealed and characterized for the first time in the sealing zone. Furthermore, cross-correlation analysis between signals associated with two cores in the same cell brought to light the existence of a spatial synchrony between neighbors up to an approximate 2 μm distance scale. Interestingly, this distance coincides with the mean neighbor-to-neighbor distance measured by SEM and RIM. Similar observations in human macrophage podosomes have been reported, showing that under a distance of 1.8 μm, podosome neighbors oscillate synchronously ^30^. A role of connecting actin filaments was suggested to be involved in controlling the synchrony between podosomes ^30, 31^. In sealing zones, the spectacularly dense actin meshwork observed between actin cores could be involved in connecting subunits and thus conducting synchronized oscillations of actin within the superstructure. In addition, our previous observations in macrophages revealed that actin oscillations corresponded to oscillations of the protrusion force generated by podosomes ^30^. Based on these observations in macrophages, we propose that in osteoclasts, synchrony of actin oscillations in actin core islets could reflect the involvement of oscillating protrusion forces in the sealing process. Actually, we also revealed the existence of intertwined islets of podosomes: some could be involved in sealing, while neighboring islets are concomitantly relaxing. A local co-regulation between actin polymerization in the core and formation of surrounding adhesion proteins was observed, as described in scattered podosomes of macrophages or dendritic cells ^27, 32^. Thus, intermittent protrusive and relaxing capacity of podosome islets could enable efficient sealing of the osteoclast plasma membrane to the bone surface and maintaining the resorption lacuna and the diffusion barrier.

This study consists in the first extensive and quantitative study of the nanoscale organization and dynamics of the sealing zone in human osteoclasts. It proposes precise localization of six major components of the sealing zone: vinculin, paxillin, talin, cortactin, filamin A and α-actinin1. Furthermore, it provides the geometric characterization of the actin core network within this unique and functional superstructure. It also allows identification of dynamic processes related to podosomes during active bone resorption. This work therefore aimed at paving the way for future studies to decipher both the ultrastructural and dynamic properties of this unique osteoclast structure dedicated to bone resorption.

## Materials and methods

### Differentiation and culture of primary monocyte-derived osteoclasts

Human monocytes were isolated from blood of healthy donors as previously described ^35^. Cells were re-suspended in cold PBS supplemented with 2 mM EDTA, 0.5% heat-inactivated fetal calf serum (FCS) at pH 7.4 and magnetically sorted with using magnetic microbeads coated with antibodies directed against CD14 (Miltenyi Biotec). Monocytes were then seeded on plastic at 2×10^6^ cells/well in six-well plates in RPMI 1640 (Invitrogen) without FCS. After 2 h at 37°C in humidified 5% CO_2_ atmosphere, the medium was replaced by RPMI containing 10% FCS, 100 ng/mL of macrophage colony-stimulating factor (M-CSF, Peprotech) and 60 ng/mL of human receptor activator of NF-κB-ligand (RANK-L, Miltenyi Biotec). The medium was then changed every third day by RPMI containing 10% FCS, 100 ng/mL of RANK-L and 25 ng/mL of M-CSF. For experiments, cells were harvested at day 10 using Accutase solution (Sigma-Aldrich) and centrifugation (1100 rpm, 10 min), and were then plated either on clean 1.5H precision glass coverslips (Marienfeld 0117640) or on bovine bone slices (Immuno Diagnostic Systems DT-1BON1000-96). Cells were left to adhere in cytokine-supplemented medium for 3 days before fixation.

### Cleaning of precision glass coverslips

Before letting cells adhere on them, 1.5H precision glass coverslips (Marienfeld) were cleaned as follows: they were placed on staining racks (Thermo Scientific 12627706) and immersed in a RBS 35 solution (Carl Roth 9238, 1/500 diluted in milliQ water) heated up to 80°C while stirred with a magnetic bar. After 15 min, coverslips were rinsed three times with milliQ water and patted dry. They were put in a dry oven set at 175°C for 120 min to sterilize. These clean coverslips were used within two weeks after RBS treatment.

### Primary antibodies

The following antibodies were used: goat anti-talin C-20 (Santa Cruz sc-7534, IF 1/50), mouse anti-vinculin clone hvin-1 (Sigma-Aldrich V9131, IF 1/50), mouse anti-paxillin (BD Biosciences 61005, IF 1/50), mouse anti-cortactin clone 4F11 (p80/85) (Sigma-Aldrich 05-180, IF 1/50), mouse anti-filamin A clone PM6/317 (Sigma-Aldrich MAB1678, IF 1/50), and mouse anti-α-actinin1 clone BM-75.2 (Sigma-Aldrich A5044, IF 1/50).

### Immunofluorescence

Osteoclasts plated on glass coverslips and bone slices for 3 days were fixed for 10 min in a 3.7% (wt/vol) paraformaldehyde (Sigma Aldrich 158127) solution containing 0.25% glutaraldehyde (Electron Microscopy Sciences 16220) in Phosphate Buffer Saline (PBS) (Fisher Scientific) at room temperature. When indicated, before fixation, cells were mechanically unroofed at 37°C using distilled water containing protease inhibitors (Roche) and 10 μg/mL phalloidin (Sigma-Aldrich P2141) at 37°C: cells were left still in this solution for 10 s, then a flow was created by flushing a dozen times so that cell dorsal membranes were ripped off. After fixation, quenching of free aldehyde groups was performed by treatment with PBS/50 mM ammonium chloride and PBS/1 mg/mL sodium borohydride. Non-unroofed cells were permeabilized for 10 min with PBS/0.3% Triton and all cells were blocked with PBS/1% BSA for 30 min. Samples were incubated with the primary antibodies for 90 min and then during 60 min with fluorescent dye conjugated-phalloidin and secondary antibodies for F-actin and proteins, respectively.

### DONALD 3D super-resolution imaging

Osteoclasts plated on cleaned glass coverslips of accurate thickness (0.170 +/-0.005 mm, Marienfeld 0117640) were unroofed and fixed as described above. Vinculin, paxillin, talin, cortactin, filamin A or α-actinin1 were stained with the corresponding primary antibody and an Alexa Fluor 647-coupled secondary antibody (Molecular Probes A21237, 1/1000) for dSTORM and podosome cores were labelled with Alexa Fluor 488-phalloidin (Molecular Probes A12379, 1/500) for epifluorescence. All dSTORM experiments were performed with the Smart-kit buffer (Abbelight, France).

3D super-localization images were acquired using an inverted IX83 microscope (Olympus) combined with SAFe module (Abbelight, France), and a TIRF module (Abbelight, France). Samples were excited with 405 nm (200mW, ERROL Laser), 488 nm (150 mW, ERROL Laser), and 640 nm (400mW, ERROL Laser) lasers in a HILO illumination (Highly Inclined and Laminated Optical sheet), and controlled *via* NEO Software (Abbelight, France). GFP / Alexa 532 / mCherry / Alexa 647 fluorescence filters (Semrock, LF-405/488/532/635-B-OFF) were used, and the objective was a 100x/1.49 N.A oil immersion objective (Olympus). All images were acquired using a sCMOS ORCA FLASH4.0 v3 (100 fps, cable camera link, Hamamatsu) camera, split on two regions of 300×300 pixels area and positioned on the focal plane of the SAFe module (2.7x magnification, optical pixel size of 108 nm). The two imaging paths are calibrated in terms of transmission efficiency to define a permanent correction factor that compensates the imperfect beam splitter. Images were collected once the density of fluorescent dye was sufficient (typically, under 1 molecule.μm^−2^). About 5000 frames were recorded to compute one image of protein, and 500000 frames were acquired to obtain one image of actin. For all recorded images, the integration time was set to 50 ms and the EMCCD gain to 150. Laser power was adapted depending on the fluorophore density.

### Analysis of DONALD 3D super-resolution images

The super-localization of molecules and drift correction were performed on raw images *via* NEO Software (Abbelight, France) to achieve a final pixel size of 15 nm. Images were grouped in batches accounting for 5% of the total frames, and were submitted to drift correction by comparison batch per batch and according to a sliding window. Eventually, z correction was performed thanks to a dedicated Python script, to account for the calibration value of the SAFe module.

The spatial organization of proteins within the sealing zone like was characterized following an adapted version of the algorithm described previously ^27^. Briefly, actin cores in-plane coordinates (x_core_,y_core_) were determined from actin epifluorescence images from the 488 nm channel after Gaussian-filtering. The angle between the podosome belt portion and the horizontal was measured by the user. Then, centered on each core, a rectangle bounding box of length 10 μm and width 200 nm was drawn and the 3D coordinates (x_i_, y_i_, z_i_) of the molecules inside this box were converted to a local r-z space (r_i_, z_i_). This operation was repeated twice for each core: once along the structure direction, and once in the transversal direction. This analysis was performed for all cores of a given cell.

To further the analysis of the spatial organization of molecules within the podosome belt, a dedicated Python script was written to extract statistical information. Each cross-section direction for each cell was treated independently, except for distributions along r-axis and along z-axis for which all molecules detected for all cells were considered. Points were sorted in classes of varying lengths, depending on their distance to the corresponding actin core and whether they were located towards the exterior or the internal part of the cell. For each r-class, median height of the distribution was estimated cell by cell. Also, the symmetry between internal and external distribution of proteins in terms of quantity was assessed by comparing the total amount of molecules on the exterior side to the total amount of points on the interior side, normalized by the total amount of points detected for each cell considered (Figure S1).

### Scanning Electron Microscopy imaging

Osteoclasts plated on bone slices for 3 days were unroofed as described above and fixed for 10 min in a 0.2M sodium cacodylate buffer (pH 7.4) containing 2% paraformaldehyde (Electron Microscopy Sciences 15710) and 2.5% glutaraldehyde (Electron Microscopy Sciences 16220), then washed with distilled water. The samples were then prepared for observation following the protocol: they were dehydrated through a graded series (25–100%) of ethanol, transferred in acetone and subjected to critical point drying with CO2 in a Leica EM CPD300, then sputter-coated with 3 nm platinum with a Leica EM MED020 evaporator, and were examined and photographed with an FEI Quanta FEG250.

### Analysis of SEM images

Actin cores were manually encircled using ImageJ oval selection tool, then the area and its center coordinates of these selected regions were extracted. A dedicated Python script was written to extract statistical information. The inter-core distances were computed using Delaunay tessellation on each image, using SciPy spatial algorithm “Delaunay” in order to create the interconnection matrix, then the distance between each pair of connected vertices was computed and stored in a matrix. In order to avoid plausible errors, the edges of the Delaunay tessellated space were excluded of the distance computations. To identify them, the SciPy spatial algorithm “ConvexHull” was applied to the coordinates, and whenever two points were identified as pertaining to the convex envelope, the distance between them was not computed. Finally, for each vertex the minimum distance to all neighbors was kept for the nearest neighbor analysis.

### RIM 2D super-resolution imaging

For live imaging, osteoclasts were transduced with GFP- or mCherry-tagged LifeAct and GFP-paxillin lentiviruses (BiVic facility, Toulouse, France) 3 days as previously described ^30^, before being harvested and plated on bovine bone slices, and were observed the day after being plated on bone slices. Bone slices were placed on a FluoroDish (WPI FD35-100) with cells facing down and immersed with RPMI without phenol red, supplemented with 10% FCS (Thermo Fisher 32404-014). During observations, samples were maintained at 37°C in humidified 5% CO2 atmosphere. Images were acquired every 12 ms for a total of 211 seconds (streams) using an inverted microscope (TEi Nikon). A fiber laser combiner with 4 fast diode lasers (Oxxius) with respective wavelengths 405 nm (LBX-405-180-CSB,) 454 nm (LBX-445-100-CSB), 488 nm (LBX-488-200-CSB), and 561 nm (LMX-561L-200-COL) are used to excite fluorophores. A corrected fiber collimator (RGBV Fiber Collimators 60FC Sukhamburg) is used to produce collimated TEM_00_ 2.2 mm diameter output beam for all wavelengths. The polarization beam is rotated with an angle of 5 degrees before hitting a X4 Beam Expander beam (GBE04-A) and produces 8.8 mm beam TEM_00_ beam. A fast spatial light phase binary modulator (QXGA fourth dimensions) is conjugated to the image plane to make speckle random illumination. The objective lens used in experiments is a 100x magnification with 1.49 numerical aperture (CFI SR APO 100XH ON 1.49 DT 0;12 NIKON). A band pass filter was used for Green Fluorescence Protein (GFP) emission (Semrock FF01-514/30-25). A motorized high speed wheel filter is used to sequentially turn the two band pass filters in 30 ms after each 200 speckle fames. A piezoelectric Z stage (Z INZERT PIEZOCONCEPT) is used for fast z stack acquisition. For triggering, the camera sCMOS is used as master, and a rolling shutter output is used to trigger binary phase sequence to the SLM. The SLM output triggers the laser when binary phase mask is stable. A script from micromanager software was written to select the number of speckles used, the temporal resolution for each frame, the depth of z stack, the step of each z stack and the number of colors used for the acquisition. For stream recording, speckle frames were acquired continuously over the whole duration of the movie. Widefield time-lapse acquisitions were also carried out with the same microscope, but using a 60x objective (CFI PLAN APO LBDA 60XH 1.4/0.13 NIKON). Images were captured every 2 seconds, and were were restored with Huygens Software (classical maximum likelihood estimation with 30 iterations and theoretical PSF). For imaging of fixed samples, osteoclasts were unroofed and fixed as described above and stained for vinculin, cortactin, filamin A, or α-actinin1 with the corresponding primary antibody and an Alexa Fluor 488-coupled secondary antibody (Cell Signaling Technology #4408, 1/500) or an Alexa Fluor 546-coupled secondary antibody (Molecular Probes A11056, 1/500). Actin cores were labelled with Texas Red-phalloidin (Molecular Probes T7471, 1/200) or Alexa Fluor 488-phalloidin (Molecular Probes A12379, 1/200) respectively. Bone slices were placed in a FluoroDish, upside down on a droplet of Vectashield mounting medium (Vector Laboratories H-1000). Samples were excited with 488 nm and 561 nm laser diodes with the same setup as for time-lapse imaging. 200 speckle images for each channel were acquired sequentially to yield z stacks of various depths.

### Analysis of RIM images of fixed samples

Reconstruction of raw images was carried out as described in ^29^.

Briefly, the method is based on decreasing the computational cost of the inversion method described in ^36^. This new method uses a variance matching process, instead of the marginal minimization methods based on the full covariance matrix of the data ^29^. The only input for the super-resolution reconstruction process is the knowledge of spatial statistics of speckle patterns limited by the OTF of the imaging system (Fourier transform of the PSF). Contrary to SIM, the exact knowledge of the illumination function is not necessary, and the protocol of the reconstruction is therefore drastically reduced. The inversion code is implemented on Matlab. The input are the excitation and collection PSF, generated with Gibson and Lanny 3D optical model implemented in the plugin PSF generator ^37^. The PSF dimension is equal to the final size reconstruction with a pixel size equal to 32.25 nm for 100x magnification. The position of the fluorophores is defined from the cover slide in each sample. The number of iterations during the variance matching process is defined by the user, mainly depending on the signal to noise ratio of the raw data.

Drift correction was performed on z stacks with ImageJ plugin Linear Stack Alignment with SIFT. To create images on which to perform further analyses, 3 to 5 z slices per acquisition stack were selected on their sharpness, depending on the quality of the original signal, and summed with ImageJ Z Project tool.

In order to characterize the actin network, actin cores were detected as local maxima using ImageJ Find Maxima tool with the threshold set as half of background intensity. The coordinates of these maxima were weighted considering all pixel intensity in a 200 nm radius, and exported in a text file. These points were positioned both on RIM reconstructed image and on its spatial derivative version created thanks to ImageJ Find Edges tool. From these locations, 8 radius profile were traced with length 1 μm and width 100 nm on both images, and intensity values along this line were extracted and stored in a text file. A dedicated Python script was then written to extract statistical data. Core radii were computed by detecting the first intensity maximum along the Find Edges profile. Intercore distances were computed thanks to weighted coordinates following the same Delaunay algorithm already described in the SEM analysis section (Figure S4).

For the analysis of two-color acquisitions, actin cores were detected following the same procedure as described for the analysis of the actin network. Then, both signals were extracted along 1.5 μm long and 100 nm wide lines drawn so that each core coordinates were placed at the middle of the line, and its orientation either followed the local curvature of the sealing zone, or was transverse. This process was repeated on the reconstructed actin image, the actin image after spatial derivation with ImageJ Find Edges tool and the protein image. All data were extracted and stored in a text file to be read by a dedicated Python script. This script computed the median core size for all data thanks to Find Edges signals, in order to establish a normalized profile length. Then, both actin- and protein-associated signals were interpolated along this new axis to yield comparable intensity profiles. Median profiles were eventually computed (Figure S5).

### Analysis of RIM live acquisitions

Reconstruction of raw images was carried out as described previously. To combine robust statistical estimation of object and temporal resolution, an interleaved reconstruction method has been made as previously proposed for SIM ^38^. 800 speckles were grouped to reconstruct one time slice, and the time step between two images corresponds to 200 speckles. Drift correction was performed on stacks with ImageJ plugin Linear Stack Alignment with SIFT, with the same parameters as for fixed samples. Actin intensity levels were normalized throughout the stack by using ImageJ Bleach Correction tool, with the correction method set to histogram matching.

To detect the single actin cores, all the time slices were summed with ImageJ Z project tool and the coordinates were extracted according to the same procedure as for fixed samples. The dynamic characteristics of actin were assessed by extracting the time-dependent signals in a circular selection or radius 100 nm around each core, and storing them in a text file. A dedicated Python script was developed to compute the distance between all core pairs in the same cell from, the Fourier spectrum associated with each signal and the Pearson cross-correlation between two signals in the same cell. Natural frequencies were identified as the frequencies associated with Fourier coefficients greater than a threshold value proportional to the median value for Fourier coefficients over the spectrum. Pearson coefficients were eventually sorted according to the distance between the core pair coordinates.

In order to obtain graphical representations of the local evolution of fluorescence over time, as shown in Figure 3, 5 and Movies3, 5, the films were processed using ImageJ as follows. First, the films were registered using the StackReg ImageJ plugin. Then, 2 sequential time points were subtracted to obtain a new film representing the rate of change of fluorescence. Finally, a Gaussian filter (2 pixel radius, i.e. 64.5 nm) and a time average on three consecutive images were applied to this film to help reduce local noise and visually highlight local variations in intensity changes.

### Statistical analysis

All box-and-whisker plots show the median, lower and upper quartiles (box) and the 10th and 90th percentiles (whiskers).

## Resource availability

All data and codes used in the analysis are available upon request to the corresponding authors.

## Acknowledgements

The authors are grateful to Myriam Ben Neji for isolation of human blood monocytes and Isabelle Fourquaux from TRI imaging facility for SEM preparation. The authors also acknowledge Anne Blangy, Alessandra Cambi, Amsha Proag and Olivier Destaing for helpful discussions. This work has been supported in part by l’Agence Nationale de la Recherche (ANR16-CE13-MechanOCs), l’Université de Toulouse, la Région Occitanie, la Fondation pour la Recherche Médicale (FRM DEQ2016 0334894), INSERM Plan Cancer and Human Frontier Science Program (RGP0035/2016).

## Contributions

MP performed and analyzed all experiments. TM and RP participated in RIM experiments. NE participated in DONALD experiments. BRM and CV participated in differentiating osteoclasts. RP and CT supervised the project. CV, CT and IMP obtained funding. MP and RP wrote the manuscript with input from the others.

## Supplementary figure legends

**Supplementary Figure 1.**
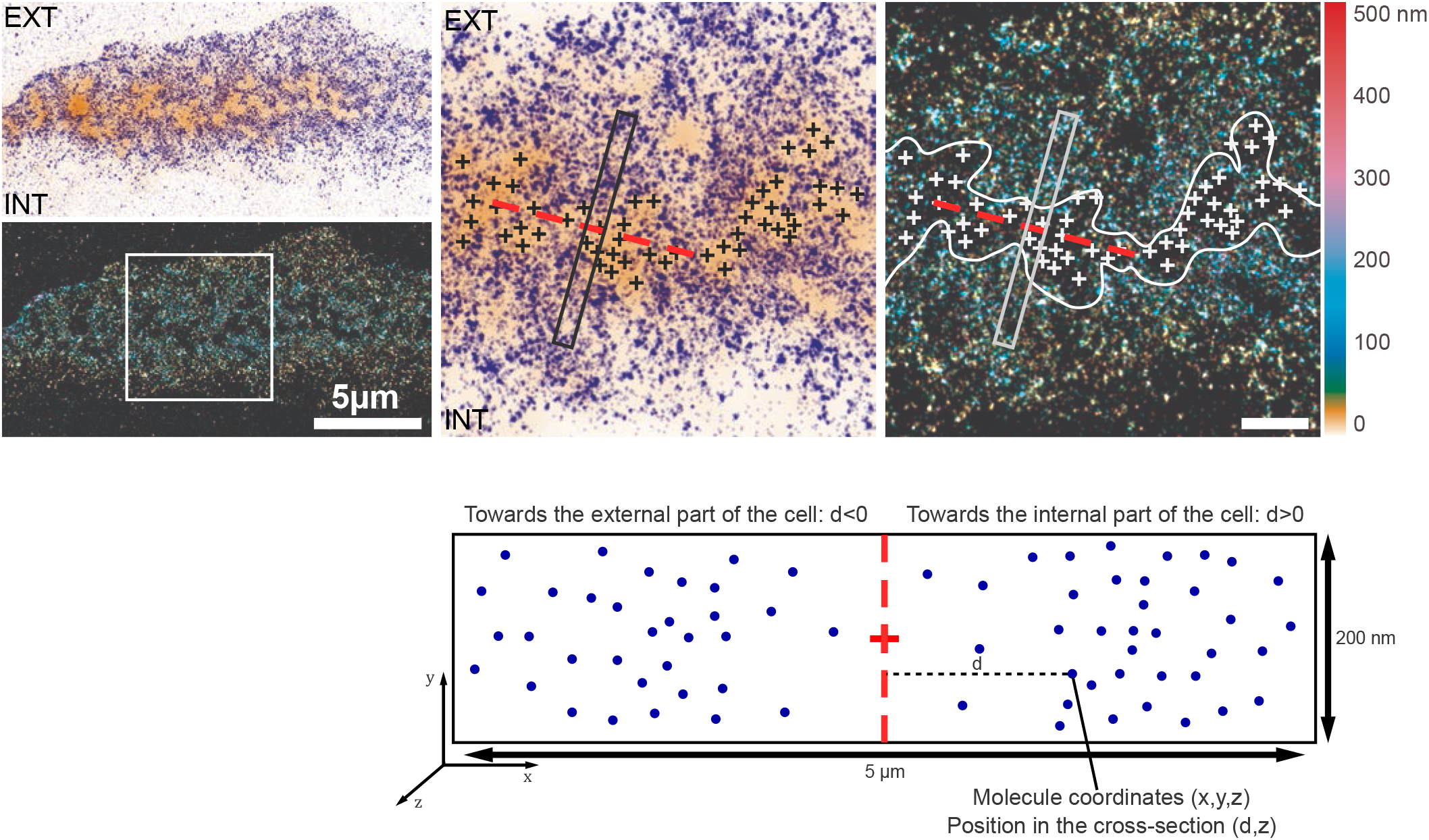
Analysis workflow for the 3D localization of proteins relative to actin cores with DONALD. When zooming in on a region of the belt, each of the actin cores (crosses) in the different clusters were located by the user. Then, the general orientation of the cluster was evaluated, and a rectangular region was delimited and centered on an actin core, transverse to the direction of the cluster. Within these cross-sections, molecules were automatically localized depending on their distance to the central axis of the cross-section and their height. Importantly, all molecules situated towards the cell edge were represented on the left, while those towards the interior of the cell are on the right in the associated charts.

**Supplementary Figure 2.**
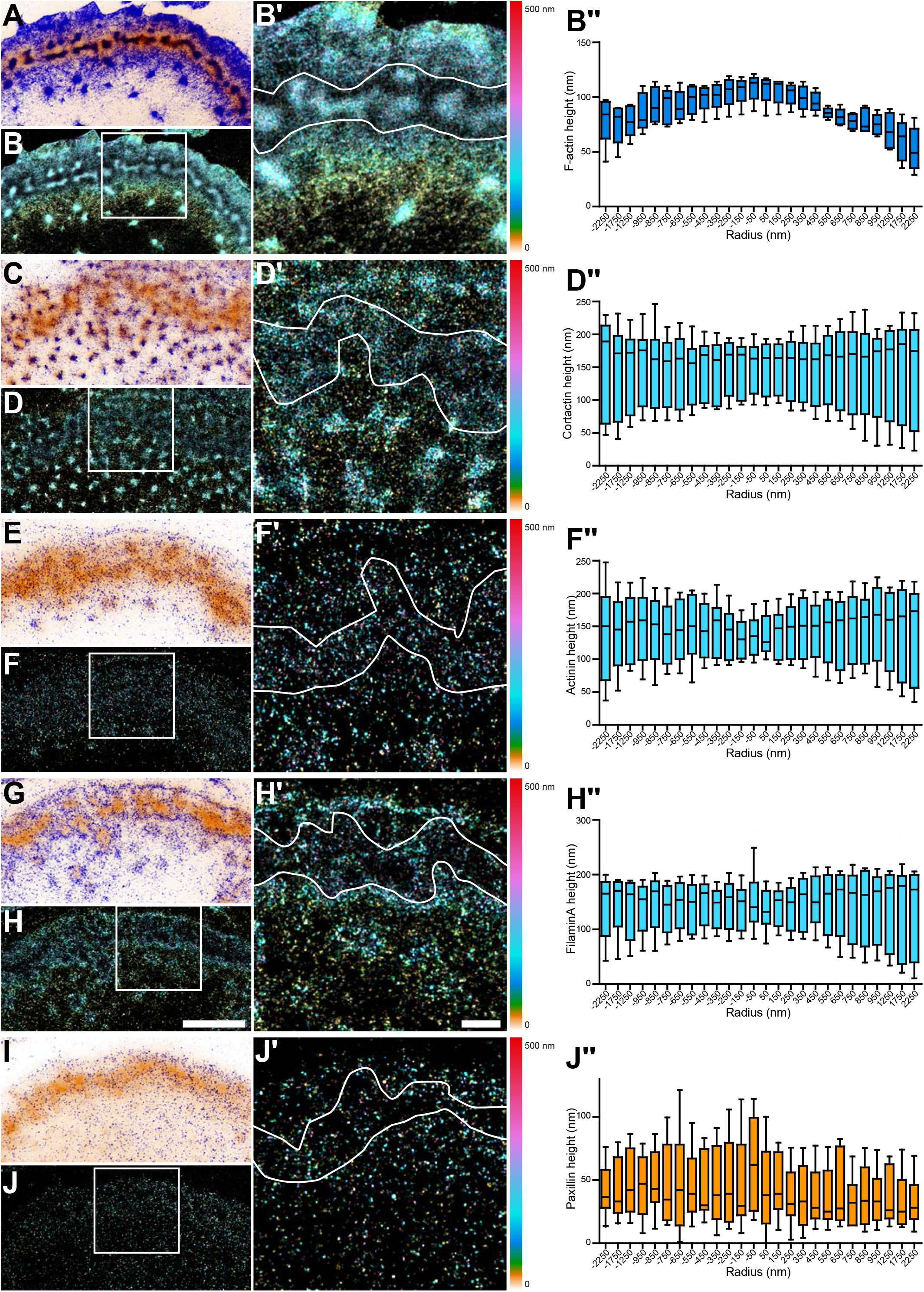
3D nanoscopy of F-actin, cortactin, α-actinin1, filamin A and paxillin in the osteoclast podosome belt. (A) Representative dSTORM images of cortactin (blue) merged with the corresponding epifluorescence images of the F-actin cores (ochre). (A’) DONALD images corresponding to (A) where the height is represented in false color (scale shown in (A’’). (A’’) Enlarged view of (A’). (A’’’) Height profiles of cortactin with respect to the distance to the center of the sealing zone. (B) Representative dSTORM images of α-actinin1 (blue) merged with the corresponding epifluorescence images of the F-actin cores (ochre). (B’) DONALD images corresponding to (B) where the height is represented in false color (scale shown in (B’’). (B’’) Enlarged view of (B’). (B’’’) Height profiles of α-actinin1 with respect to the distance to the center of the sealing zone. (C) Representative dSTORM images of filamin A (blue) merged with the corresponding epifluorescence images of the F-actin cores (ochre). (C’) DONALD images corresponding to (C) where the height is represented in false color (scale shown in (C’’). (H’’) Enlarged view of (C’). (C’’’) Height profiles of filamin A with respect to the distance to the center of the sealing zone. (D) Representative dSTORM images of paxillin (blue) merged with the corresponding epifluorescence images of the F-actin cores (ochre). (D’) DONALD images corresponding to (D) where the height is represented in false color (scale shown in (D’’). (D’’) Enlarged view of (D’). (D’’’) Height profiles of paxillin with respect to the distance to the center of the sealing zone. Scale bars: 5 μm (A’, B’, C’, D’), 1 μm (A”, B”, C”, D”).

**Supplementary Figure 3.**
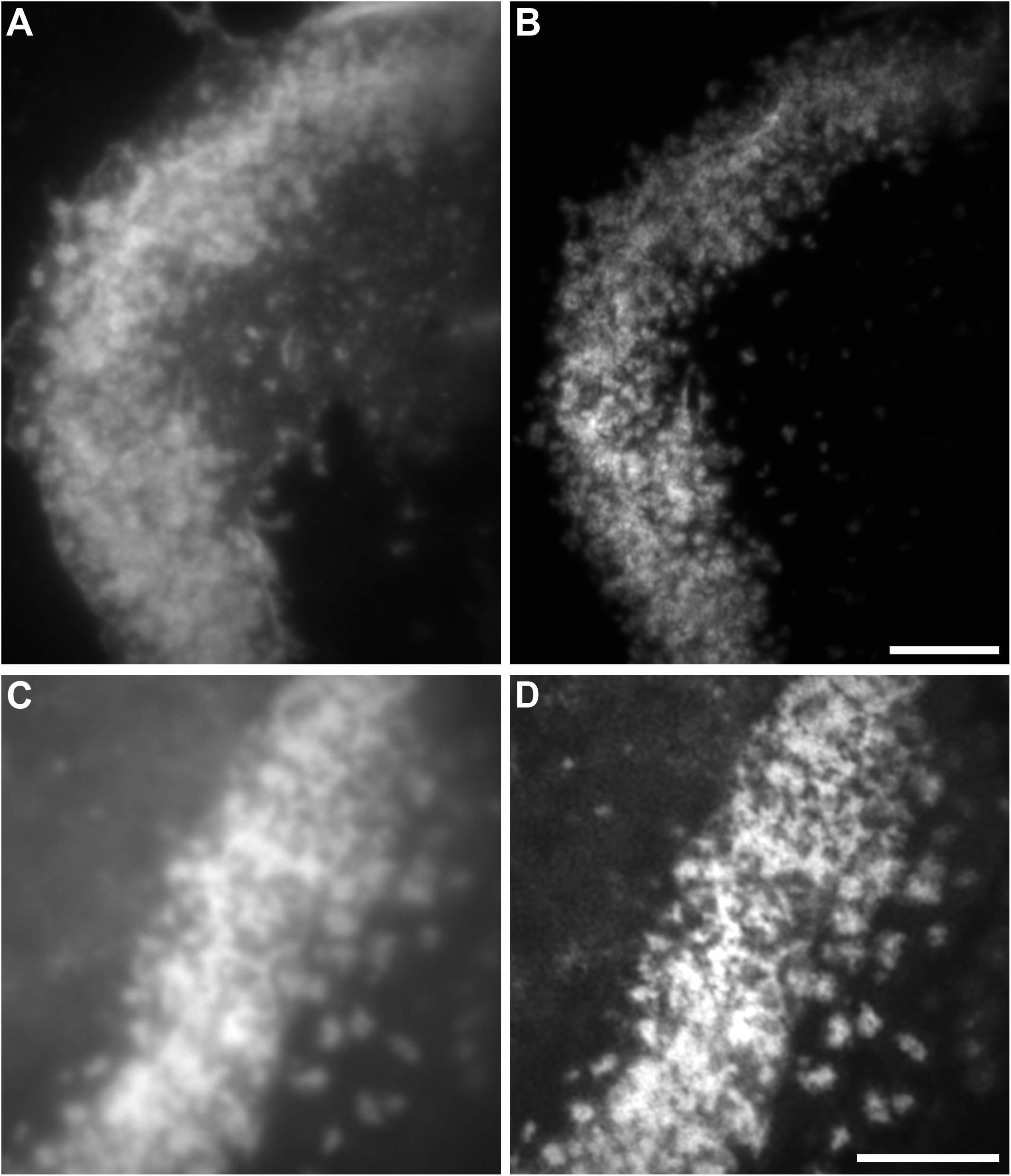
Comparison between epi-fluorescence and RIM super-resolution microscopy. (A) Widefield image of the sealing zone stained for F-actin already shown in Figure 1C. (B) RIM image of the sealing zone shown in Figure 1C. (C) Widefield image of the sealing zone stained for F-actin already shown in Figure 2B. (D) RIM image of the sealing zone shown in Figure 2B.

**Supplementary Figure 4.**
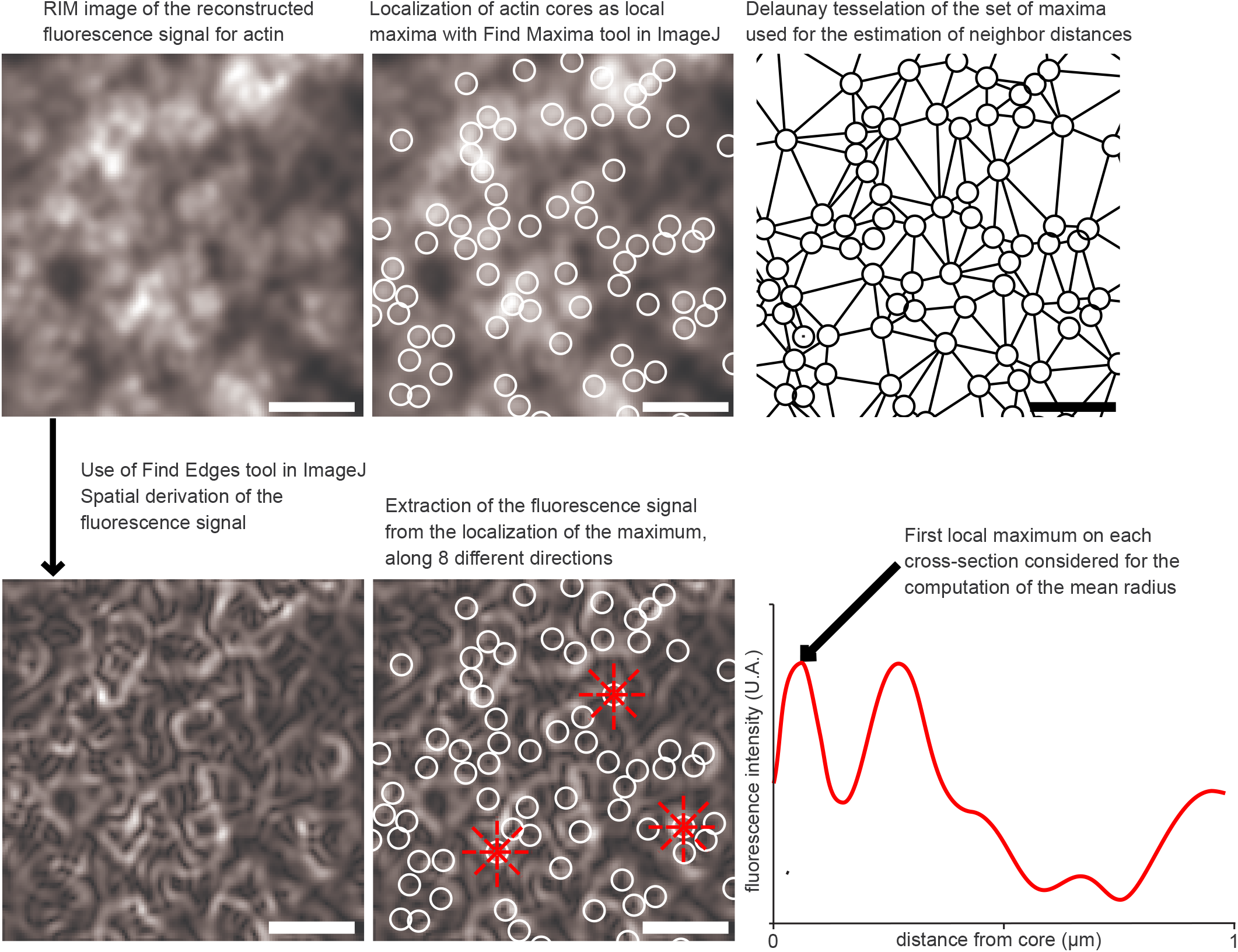
Analysis workflow for the geometric characterization of actin cores with RIM technique. When zooming in on a region of a sealing zone, it is observed that actin signal displays numerous local maxima that were localized. The corresponding coordinates were used to estimate the distance between two neighboring maxima, based on Delaunay’s tessellation. Then, in order to assess the size of the actin dots, a spatial derivation of the signal was applied and the derived signal was extracted along 8 radii from the center of each dot. On these curves, the first local maximum corresponded to the edge of the actin spot and helped me compute the mean radius for all of them.

**Supplementary Figure 5.**
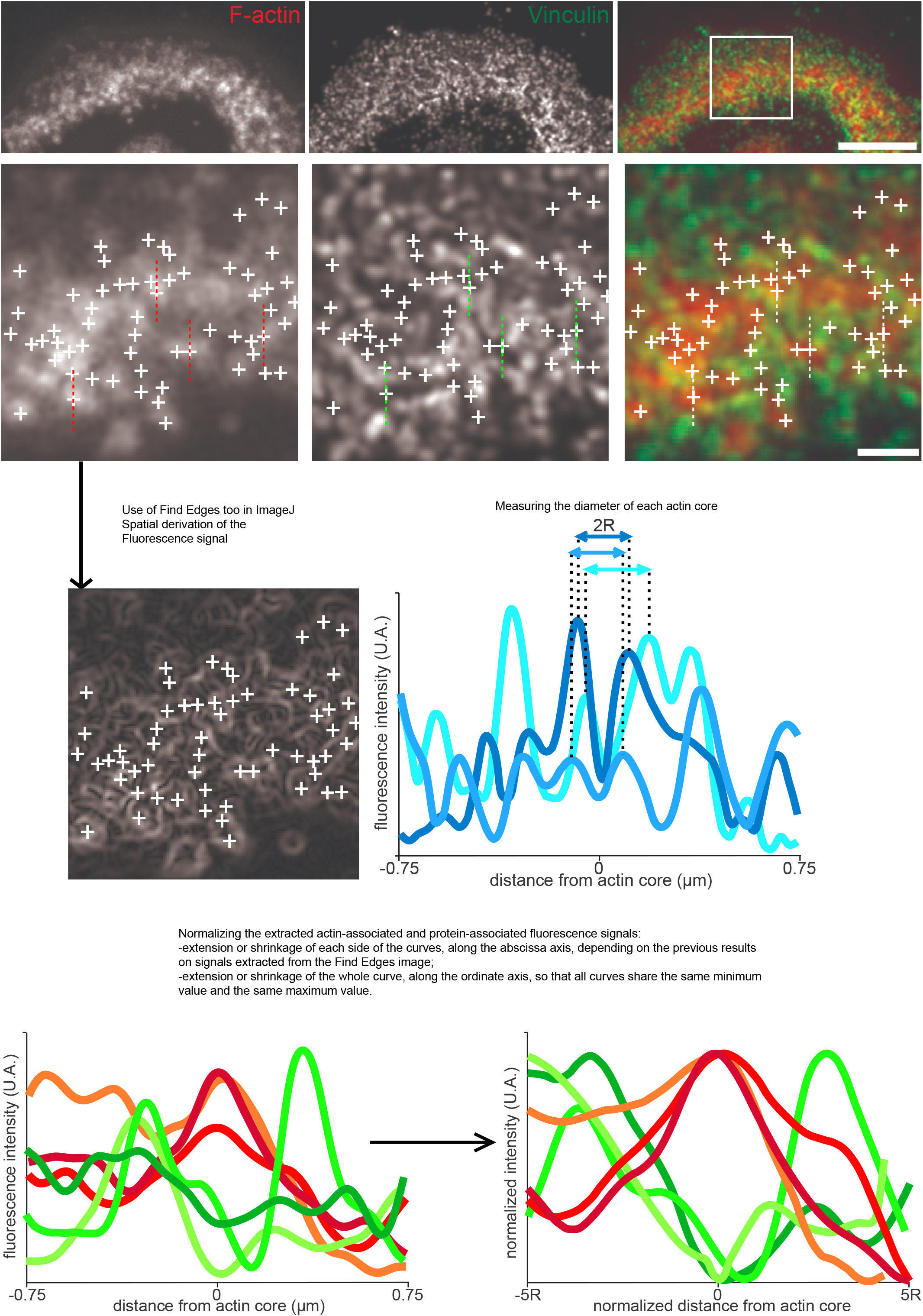
Analysis workflow for the 2D localization of proteins relative to actin cores with RIM technique. Similarly to the evaluation of the size of actin cores, the first step was to extract the coordinates of the cores thanks to the actin staining image. And in the same way, data was also extracted in order to assess the diameter of each core. Then, the local orientation of the sealing zone was evaluated by the user, in order to extract both actin and protein associated signals in the transverse direction. But these curves are not comparable as such, so thanks to the size-related data a normalization protocol was performed on the signals so that they were ready for statistical analysis.

**Supplementary Figure 6.**
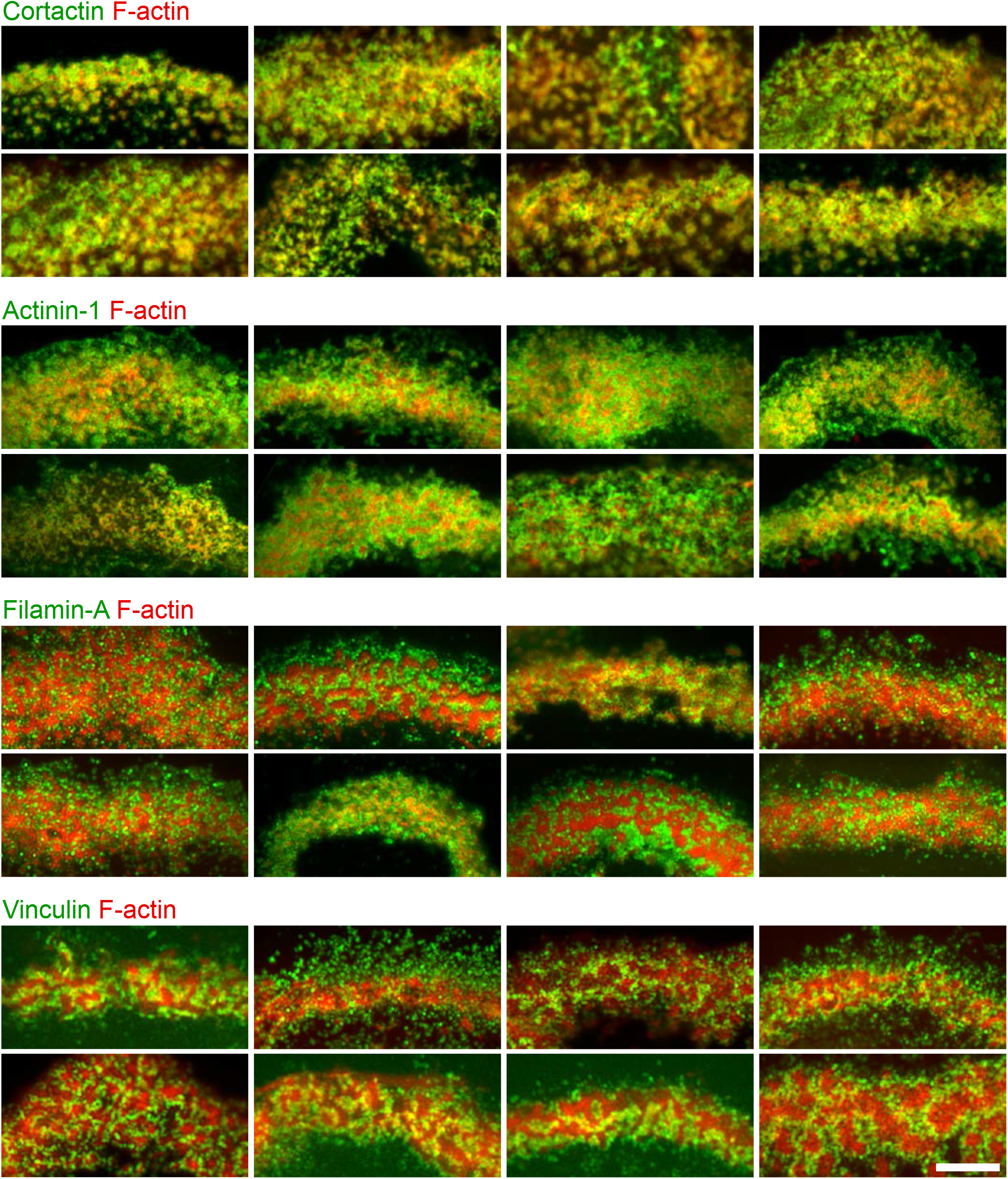
Localization in the sealing zone of cortactin, α-actinin 1, filamin A and vinculin. Gallery of immunofluorescence images of sealing zones co-stained for F-actin (red) and cortactin, α-actinin 1, filamin A or vinculin (green). Scale bar: 5 μm.

**Supplementary Figure 7.**
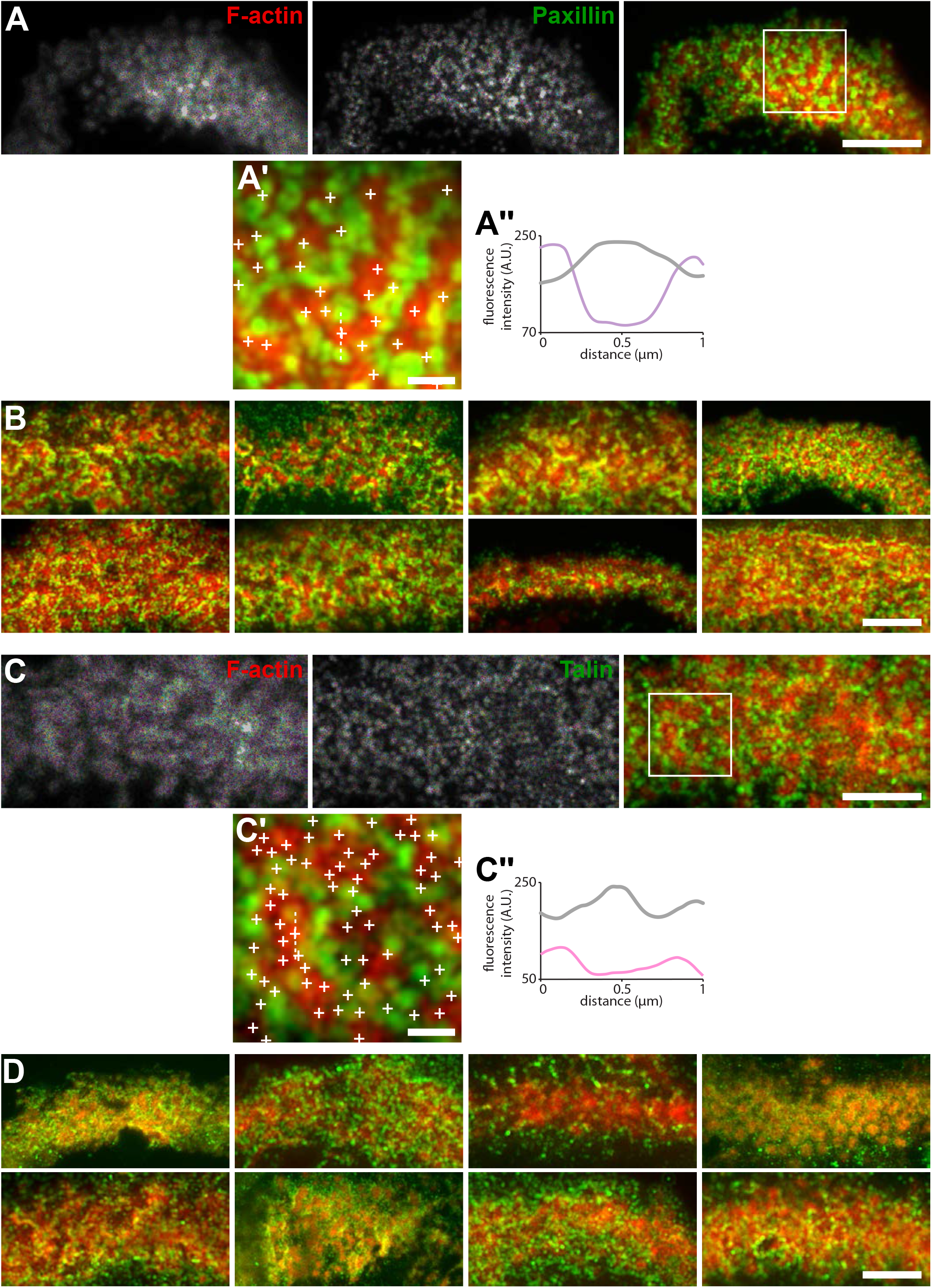
Localization in the sealing zone of paxillin and talin. (A) Representative immunofluorescence images of sealing zones co-stained for F-actin (red) and paxillin (green). (A’) Enlarged view of (A). (A’’) Intensity profiles along the dotted lines marked in (A’). (B) Gallery of immunofluorescence images of sealing zones co-stained for F-actin (red) and paxillin (green). (C) Representative immunofluorescence images of sealing zones co-stained for F-actin (red) and talin (green). (A’) Enlarged view of (A). (A’’) Intensity profiles along the dotted lines marked in (A’). (D) Gallery of immunofluorescence images of sealing zones co-stained for F-actin (red) and talin (green). Scale bars: 5 μm (A, B, C, D), 1 μm (A’, C’).

**Movie 1. Z-stack by RIM of a human osteoclast adhering on bone and stained for F-actin.**

The movie shows a stack of optical sections acquired by RIM microscopy at 200-nm intervals. Color-coded for height using a rainbow scale.

**Movie 2. Deconvolution time-series of a sealing zone over 30 min.**

Time-series movie of a human osteoclast expressing Lifeact-GFP and adhering on bone. The video was acquired by wide-field fluorescence microscopy at 2-s intervals during 30 min and deconvoluted.

**Movie 3. RIM time-series and rate of fluorescent change of the F-actin content of a sealing zone.**

Reconstruction at 2.4-s intervals during 160 s of a time-series acquired by RIM microscopy. Left panel: RIM images of a sealing zone stained for F-actin with Lifeact-GFP. Right panel: images of the rate of fluorescence change. Orange stands for a positive gradient, *i.e*. local actin polymerization, and blue represents a negative gradient, *i.e*. local actin depolymerization.

**Movie 4. Dynamics of F-actin and Paxillin in a sealing zone.**

Time-series movie by RIM microscopy of an osteoclast on bone expressing Lifeact-mCh and Paxillin-GFP, reconstructed at 9-s intervals during 160 s.

**Movie 5. RIM time-series and rate of fluorescent change of the F-actin content of a sealing zone, relative to the location of Paxillin.**

Left panel: time-series movie by RIM microscopy of an osteoclast on bone expressing Lifeact-mCh and Paxillin-GFP. A single RIM image of Paxillin-GFP at the starting point of the movie was reconstructed and superimposed on a time series of Lifeact-GFP reconstructed at 2.4-s intervals during 88 s. Right panel: images of the rate of fluorescence change of Lifeact-GFP.

